# Developmental Synchrony of Retinal Waves, Apoptosis, and Angiogenesis in Postnatal Retina

**DOI:** 10.64898/2026.03.19.712402

**Authors:** Michael A. Savage, Cori Bertram, Jean de Montigny, Courtney A. Thorne, Rachel Queen, Majlinda Lako, Gerrit Hilgen, Evelyne Sernagor

## Abstract

Postnatal mouse retinal development is a multi-faceted process involving the coordinated interaction of spontaneous neural activity as retinal waves, vascular plexus growth, and programmed cell death. While these processes are known to interact at a coarse scale, the specific mechanisms integrating them have remained elusive. Using large-scale, widefield calcium imaging, high-density multielectrode array recordings, single cell RNA-seq, and immunohistochemistry, we characterise a tightly aligned centrifugal expansion pattern during retinal development. This pattern is common to stage II retinal wave onsets, vascular development, Heme oxygenase-1 (Hmox1) expressing microglia, apoptotic cell markers, and a novel set of auto-fluorescent cluster complexes (ACCs) identified in this study. Apoptotic cells are known to upregulate functional Pannexin1 (PANX-1) hemichannels. These voltage-gated channels release purinergic molecules which act as “eat me” signals to neighbouring microglia. PANX-1 hemichannel blockade with the drug probenecid results in a profound decrease in spontaneous wave frequency and strength, suggesting that retinal waves are indeed triggered by these apoptotic cells.

Taken together, our observations suggest that spontaneous waves are initially triggered in hotspots by hyperactive apoptotic RGCs in unvascularised retinal areas. These apoptotic cells release purinergic molecules via PANX-1 hemichannels, leading to wave generation. This hyperactivity leads to local hypoxic conditions, which, coupled with high extracellular ATP concentrations, promotes angiogenesis. Once blood vessels reach a particular hotspot, ATP release activates Hmox1 positive microglia, which engulf the dying RGCs, creating the auto-fluorescent clusters. Herein, we present a unified mechanism linking causally linking early neural activity, programmed cell death and angiogenesis in the mammalian retina.

## Introduction

Spontaneous neural activity is a hallmark of developing neural tissue (O’Donovan, 1999). This is ubiquitous across species and present in both the peripheral and central nervous systems. In the retina, this takes the form of waves of electrical activity that sweep across the retinal ganglion cell (RGC) layer. It occurs during very precise and restricted temporal windows, overlapping with other important developmental events such programmed cell death, neurite outgrowth, synapse formation and angiogenesis. This research focuses on the first postnatal week at which time stage II retinal waves are present. They are mediated by cholinergic synaptic transmission through starburst amacrine cells (SACs) and last until postnatal day (P)10 (Bansal et al., 2000; Feller et al., 1996; Sernagor & Grzywacz, 1996, 1999; Wong et al., 1998; Z. J. Zhou & Zhao, 2000). While the role of spontaneous activity in refining network connectivity is well-documented, the endogenous factors that initiate and spatially constrain these waves, particularly their link to non-neuronal maturation, remain largely unexplored.

During the first postnatal week, the mouse retina undergoes angiogenesis, with the superficial vascular plexus extending from the optic nerve head at P0 towards the peripheral retina by P7 (Paredes et al., 2018). Local hypoxia and gradients in metabolic demand drive the outward expansion of the vascular plexus (Chan-Ling et al., 1995; Gariano & Gardner, 2005). Stage II retinal waves act as this metabolic driver with activity increasing in size and frequency from P2 before declining sharply at P7 (Maccione et al., 2014; Weiner et al., 2019).

Microglia have been shown to have an intimate connection to the progression of vascular development in the retina. They localise to the endothelial tip cells and interact with the developing vasculature endothelium (Liu et al., 2026). A specific subgroup of microglia expressing the *Homx1* gene has been shown to display an annular pattern of distribution around the growing vascular plexus edge (Martineau et al., 2026). Depleting microglia leads to a significant decrease in expansion and density of retinal blood vessels, and reintroduction of microglia results in reversal and recovery of the superficial vascular plexus (SVP) (Checchin et al., 2006).

Developmental apoptosis in the murine retina follows a centrifugal wave pattern mirroring both vascular growth and the microglial annulus. These apoptotic waves peak during the first postnatal week, after which they decline by P7 (Anderson et al., 2019; Farah & Easter, 2005). This process is functionally linked to remodelling via signalling molecules such as ATP, which is released through PANX-1 hemichannels and acts as a dual-purpose driver. It triggers angiogenesis (Umaru et al., 2015; Y. T. Zhou et al., 2022) while simultaneously serving as an ’eat-me’ signal that transforms quiescent, ramified microglia into the amoeboid, phagocytic state required to engulf dying cells (Chekeni et al., 2010).

While retinal wave progression, developmental apoptosis, and angiogenesis occur concurrently in the postnatal mouse retina, existing literature lacks a cohesive framework that accounts for their interdependence. Despite their overlapping timelines, these processes are frequently studied in isolation, leaving the functional links between neural activity, cell death, and vascularisation largely unexplored.

In this study, we identify a novel population of auto-fluorescent cluster complexes (ACCs) that form an annulus beneath the leading edge of the superficial vascular plexus (SVP) during the P0–P7 developmental window. We characterise the genetic makeup of these ACCs with single cell (sc) RNA-Seq which proves they are complexes of microglia, dying RGCs. This interaction is further bolstered by immunohistochemical profiling which shows that microglia are attracted to the ACCs’ local area and that a specific *Homx1* expressing subtype contains dying RGC fragments in their intracellular vesicles. We then show that purinergic signalling through highly expressed apoptotically associated PANX-1 hemichannels is both necessary for proper microglial activation, ACC formation, and stage II retinal wave generation. Together, these results provide a unified framework for the timely vascularisation of the postnatal retina.

## Results

### Centrifugal Expansion Pattern is Shared Between ACCs and Vascular Development in Postnatal Mouse Retina

The hypothesis that retinal waves, apoptosis, and angiogenesis are synchronised in the developing retina emerged from the discovery of sparse ACCs, which form a distinct, tight annulus around the optic nerve head (ONH) at P2–3 (Figure 1A). These ACCs appear sequentially in eccentricity outward from P2, disappearing at the retinal periphery by P7. This distribution precisely mirrors the expansion of the SVP, which extends from the ONH toward the periphery between P7 and P9 (Figure 1A, B, G). The ACCs exhibit broad-spectrum fluorescence across multiple wavelengths (Figure 1C) and are situated in close proximity to, but do not overlap with, starburst amacrine cells (SACs; Figure 1D). These cells are localised exclusively within the ganglion cell layer (GCL) and are entirely absent from the inner nuclear layer (INL; Figure 1F). Notably, the ACCs consistently trail the leading edge of the vascular front across all developmental stages, with peak excentric positional change observed between P2 and P3, suggesting a mechanistic role in SVP guidance (Figure 1G). Quantitative analysis further reveals significantly elevated blood vessel branching in ACC-associated regions compared to equivalent eccentricities where ACCs are absent (Figure 1E). To characterise the molecular identity of these cells and their potential role in synchronising retinal waves with angiogenesis, we performed single-cell RNA sequencing (scRNA-seq).

**Figure 1:**
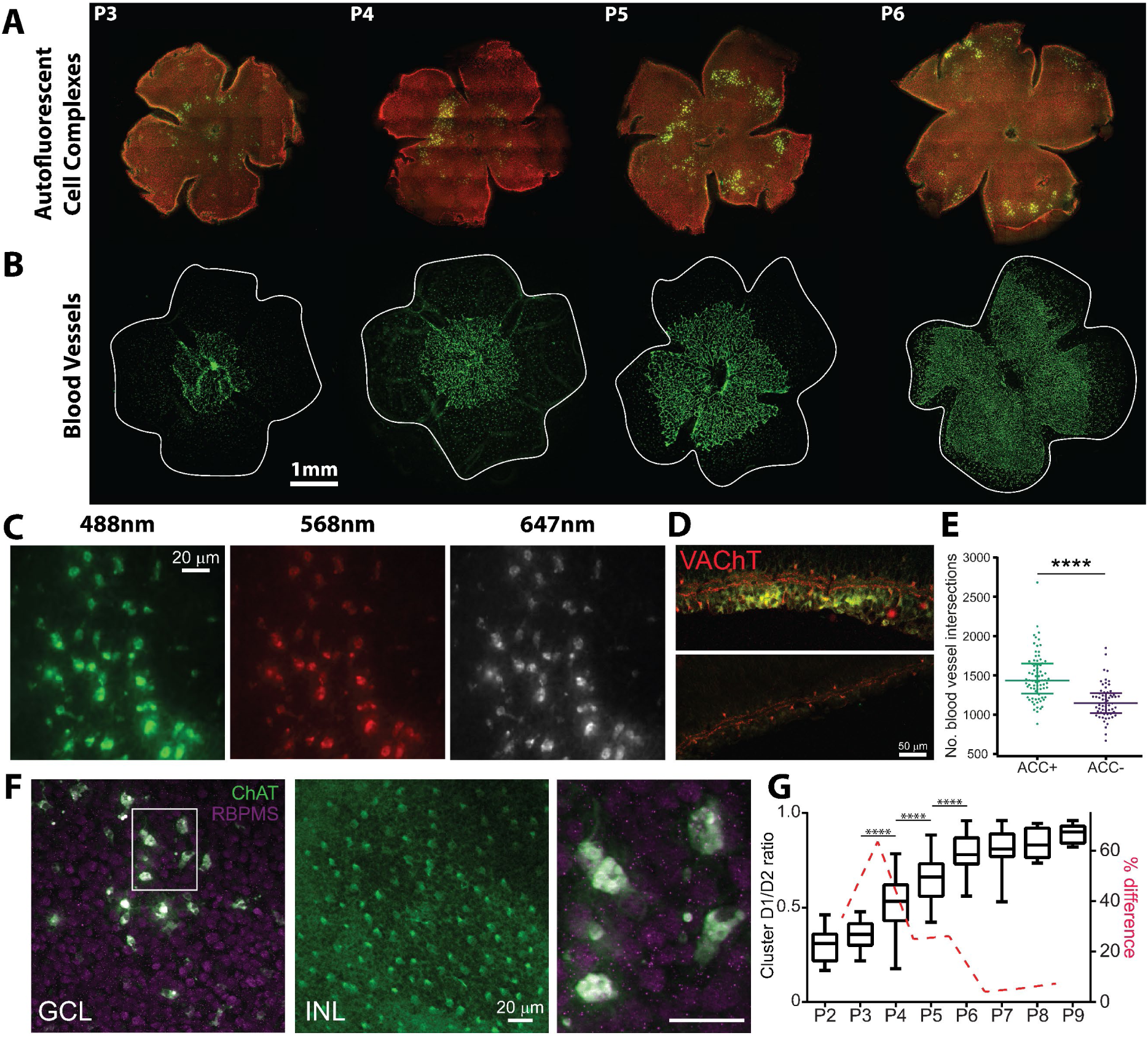
Auto-fluorescent Cluster Complexes (ACCs) display centrifugal development pattern during the first postnatal week in C57/BL6 and align with vasculature development. **A.** Mouse retinal wholemounts displaying ACCs (green) across P3-P6. **B.** Mouse retinal wholemounts (P3-P6) displaying SVP (labelled with Isolectin B4 (IB4)). Scale bar is equal to 1mm. **C.** Zoom into one auto-fluorescent cluster complex (P5) visualised at three different wavelengths without any additional immunolabeling, indicating intrinsic auto-fluorescent signals. **D.** Retinal sections showing VAChT (red) expression in a cluster area (upper panel, P4 retina) and in an area devoid of clusters (lower panel, P5 retina). **E.** Scatter plot of blood vessel branch intersections (Sholl analysis). There are significantly more branches in ACC positive areas (ACC +, green) than in ACC negative areas (ACC -, purple). Median with interquartile range. **F.** P5 cluster viewed at the GCL level and at the inner nuclear layer (INL) level in a whole-mount. Green: ChAT; magenta: RBPMS. The cluster cells are visible only at the GCL level and are much larger than SACs or RGCs (in red). At the INL level, there are only SACs and no ACCs. Right side of panel is magnified version of box on left. **G.** Box plot showing developmental changes in ACCs D1/D2 ratios. Each box illustrates the median (horizontal line) and interquartile range, with minimum and maximum values (whiskers). Asterisks indicate significant changes between consecutive days (One-way ANOVA with post-hoc Tukey test). The red dotted line illustrates the percentage difference in values between consecutive days, showing peak difference between P3 and P4 and no further changes from P6 onwards. ****: P<0.0001, Mann Whitney 2-tailed test.

### Single-Cell RNA-Seq Reveals Auto-fluorescent Cell Complexes as Microglia and Dying RGC Aggregates in the Postnatal Mouse Retina

ACCs from P5 dissociated retinas were enriched using flow activated cells sorting (FACS) and subjected to scRNA-Seq, which revealed the presence of five distinct clusters (Figure 2A). P5 was chosen as pilot data suggested that timepoint had the largest fraction of ACCs. Following FACS, the cell suspension was centrifuged to meet the input requirements of the 10x Genomics microfluidic platform, resulting in an estimated ∼50% reduction in yield. Haemocytometer quantification indicated that 9,405 viable single cells were loaded onto the 10x chip. Given an expected capture efficiency of ∼50%, approximately 5,000 FACS-purified cells were recovered for scRNA-seq library preparation. Libraries were generated and sequenced to a depth of 50,000 reads per cell using Illumina technology.

**Figure 2:**
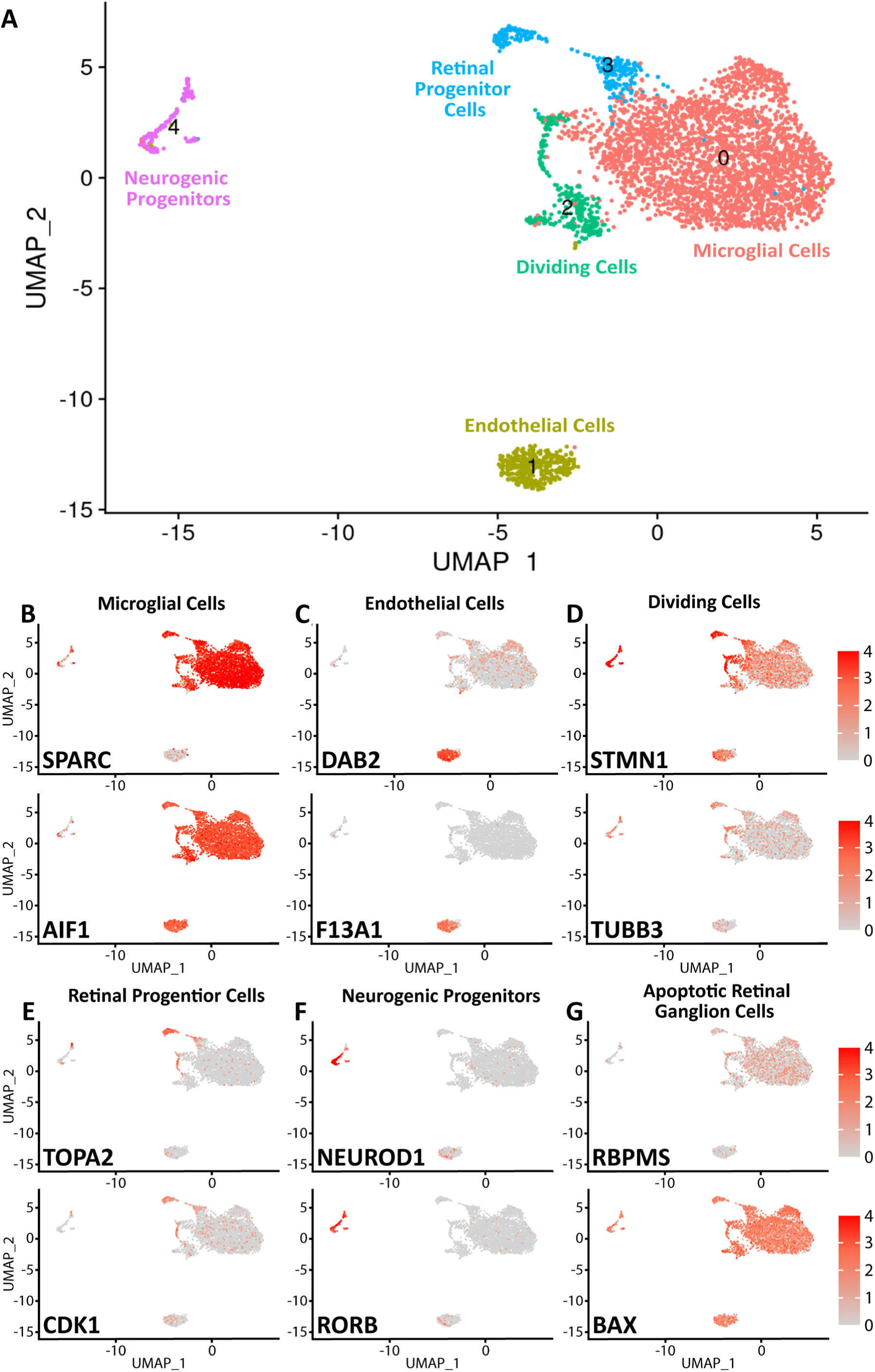
Genetic analysis of ACCs suggests they are complexes of microglia and dying RGCs. **A.** UMAP plot of scRNA-Seq data derived from ACCs taken from P5 retinas. Each cluster was identified based on the expression of retinal specific cell markers. Highly expressed markers for each cluster are shown in subsequent panels. **B.** Expression of microglia cell markers *SPARC* and *AIF1*. **C.** Expression of endothelial cell markers *DAB2* and *F13A1*. **D.** Expression of proliferating cell markers *STMN1* and *TUBB3*. **E.** Expression of RPC markers *TOPA2* and *CDK1*. **F.** Expression of neurogenic progenitor cell markers *NEUROD1* and *RORB*. **G.** Expression of retinal ganglion cell markers *RPBMS* and *BAX*.

Unsupervised clustering and UMAP visualisation identified five distinct cell populations (Figure 2A). Cluster 0 was identified as microglia, exhibiting high expression of canonical markers including *Aif1*, *Sparc*, *C1qb*, *Sall1*, *P2ry12*, *Cd68*, *Dock10*, and *Cx3cr1* (Figure 2B). Cluster 1 comprised endothelial cells, as evidenced by the expression of *F13a1* and *Dab2* (Figure 2C). Clusters 2 and 3 represented cycling populations: Cluster 2 was characterised by microtubule dynamics genes (*Stmn1*, *Tubb3*; Figure 2D), while Cluster 3 consisted of retinal progenitor cells (RPCs) expressing proliferation markers such as *Top2a* and *Cdk1* (Figure 2E). Cluster 4 was identified as neurogenic progenitors based on the expression of *Rorb* and *Neurod1* (Figure 2F).

Notably, the whole dataset exhibited high levels of the apoptotic marker *Bax* and the RGC-specific marker *Rbpms* (Figure 2G), suggesting a transcriptomic signature for ACCs consistent with RGC apoptosis and engulfment by microglia. A comprehensive list of cluster-specific markers is provided in Supplementary Table A.

### Microglia Are Attracted to ACCs in the Postnatal Mouse Retina

Given that scRNA-Seq analysis identified a prominent microglial component within ACCs, we next examined whether microglial distribution in situ was spatially coupled to the presence of ACCs. Immunolabelling for the microglial marker *Iba-1* demonstrated strong co-localisation between microglial cells and ACCs (Figure 3A). Quantitative analysis confirmed that microglial density was significantly elevated in ACC-associated regions compared with matched retinal areas lacking clusters (p = 0.0138). Across retinas, microglial density significantly increased when associated with ACCs (Figure 3B, p=0.0318), and there was a strong positive correlation observed between microglial cell density and ACC cell density (Figure 3C, p < 0.0001).

**Figure 3:**
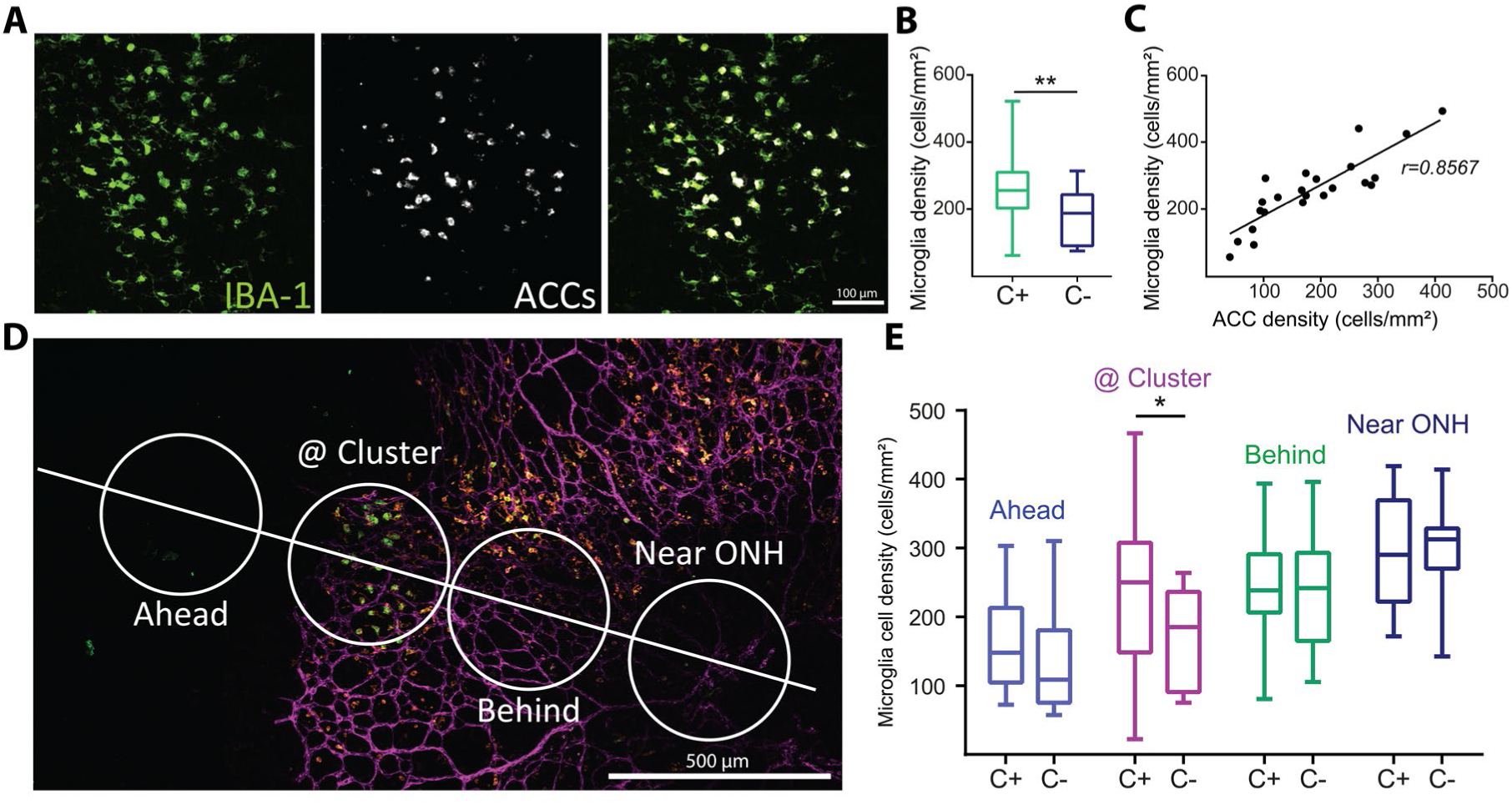
Microglia associate with the ACCs. **A.** Representative micrograph of ACCs (white) surrounded by microglial cells labelled with iba-1 (green) Scale = 100µm. **B.** Box plot (median and interquartile ranges) comparing microglia density within cluster areas (C+) versus cluster negative areas (C-). P<0.0138. Mann Whitney two-tailed test. **C.** Strong positive correlation between the densities of microglial cells and auto-fluorescent cluster cells (p<0.0001). **D.** Method for measuring microglia density at different retinal eccentricities, with measures taken near the ONH, between the ONH and clusters (behind) at clusters and ahead of clusters, further in the periphery, in the unvascularized part of the retina. The micrograph shows blood vessels lableled with IB4 (magenta), microglia labelled with iba-1 (orange) and ACCs (white) Scale bar = 500µm. **E.** Box plot (with medians and interquartile ranges) showing that the density of microglia is significantly higher within clusters (p<0.01, Mann Whitney, two-tailed test).

Because microglial density varies as a function of retinal eccentricity during postnatal development, we assessed whether enrichment at ACC sites could be explained by underlying radial gradients. Regions of interest were sampled at four progressively increasing eccentricities from the optic nerve head (ONH), with paired regions defined as ACC-positive (ACC+) or ACC-negative (ACC−) (Figure 3D). Consistent with developmental migration patterns, microglial density increased closer to the ONH in all samples (Figure 3E). However, when eccentricity was controlled, microglial enrichment was restricted to regions directly co-localised with ACCs. Only regions positioned at the same radial distance as the ACC annulus exhibited significantly elevated microglial density relative to matched ACC− regions (Figure 3E, “@ Cluster”, p<0.01). Areas located either peripheral (“ahead”) or central (“behind”) relative to ACC positions did not show a comparable increase (Figure 3E).

### Hmox1-positive Microglia Target Apoptotic RGCs through a PANX-1-Purinergic Signalling Axis

To investigate these findings further, we specifically interrogated a subset of *Hmox1*-expressing microglia. This population has been previously implicated in vascular development and remodelling, particularly in the context of retinal maturation (Martineau et al., 2026). This subset of microglia exhibits a similar annular expansion pattern during the first postnatal week to that of the ACCs and SVP (Figure 4A). These microglia straddle the vascular margin, extending processes both ahead of and behind the vessels. In these regions, they maintain direct contact with ACCs and exhibit clear evidence of engulfment, with ACC material localised within intracellular vesicles (Figure 4C).

**Figure 4:**
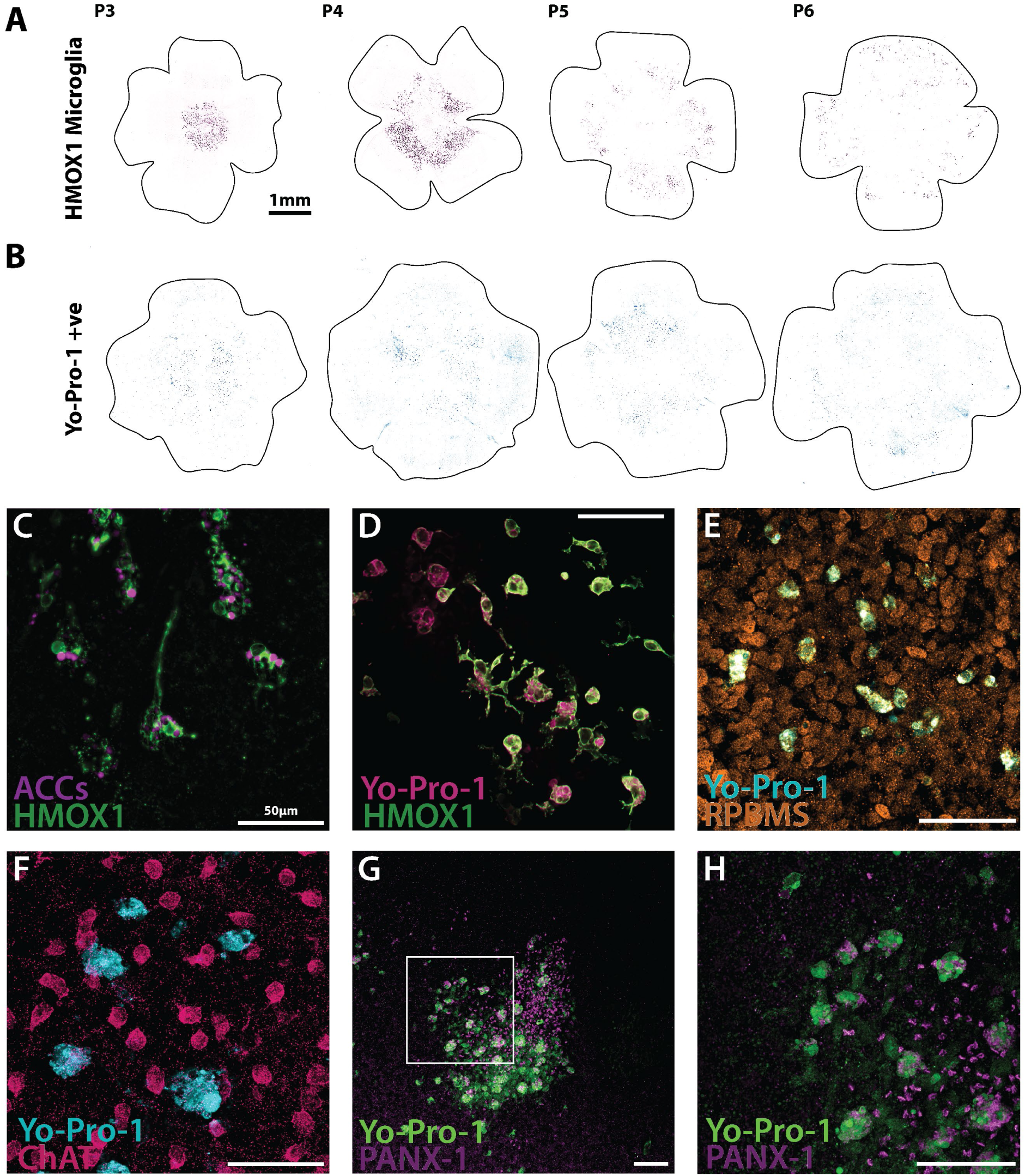
ACCs are composed of dying RGCs engulfed by Hmox1 expressing microglia. **A.** Mouse retinal wholemounts displaying a subtype of microglia labelled by Hmox1 which sit as an annulus astride the ACCs locations and the SVP. **B.** Mouse retinal wholemounts displaying apoptotic cells labelled with YO-PRO-1 in the periphery and overlapping with the microglia annulus. Scale bar 1mm. **C.** Example Hmox-1 positive microglia with ACCs present inside intracellular vacuoles responsible for breaking down and storing apoptotic cells. **D.** Apoptotic cells labelled with YO-PRO-1 are surrounded by Hmox-1 ramified microglia which become activated and engulf dying cells storing them in intracellular vacuoles similar to AAC storage. **E.** Apoptotic cells labelled with YO-PRO-1 show complete overlap with RGC marker RPBMS. **F.** Apoptotic cells labelled with YO-PRO-1 show no colocalisation with SAC marker ChAT. **G.** PANX-1 hemichannels location tightly overlaps with a cluster of YO-PRO-1 positive cells. **H.** Higher magnification inset of G displaying the tight association between YO-PRO-1 positive cells and PANX-1 hemichannels. Scale bar 50µm.

Microglial activation is frequently driven by the sensing of environmental purinergic molecules. Specifically, apoptotic cells release ATP via voltage-gated Pannexin-1 (PANX-1) hemi-channels, which is sensed by microglial P2Y12 receptors. To determine if this signalling axis contributes to ACC formation, we utilised the apoptotic marker YO-PRO-1, which selectively enters cells through open PANX-1 channels. Wholemount staining revealed a broad centro-peripheral gradient of apoptosis; however, this apoptotic annulus was positioned more peripherally than the ACCs, SVP, and *Hmox1*-positive microglia (Figure 4B). This suggests that dying RGCs are apparent across the wider periphery of the retina which then are contacted by the time lagging annulus of *Hmox1*-positive microglia.

High-resolution immunohistochemistry further demonstrated that these apoptotic cells localise within the same intracellular vesicles as the ACCs (Figure 4C, D). Co-staining with Rbpms and ChAT confirmed the identity of these YO-PRO-1-positive cells as RGCs rather than SACs (Figure 4E, F). Finally, we observed a precise spatial alignment between PANX1 expression and YO-PRO-1-positive cells, which together form an ACC-like organisation in the developing retina (Figure 4G, H). Together this suggests that RGCs initiate apoptotic mechanisms which then attract the attention of phagocytotic *Hmox1*-positive microglia which then move to engulf the cells.

### Purinergic Signalling Modulation Drastically Affects Hmox1 Microglia Activation and ACC Formation

Hmox1 microglia activation is mediated via purinergic signalling through PANX-1 hemichannels in apoptotic cells, acting as a ‘find me, eat me’ signal. This ATP sensing is carried out by P2Y12 receptors on the microglial cell surface. To investigate and verify this fully, we examined the effects of blocking either ATP release through the PANX-1 channels with probenecid or ATP sensing from microglia via P2Y12 receptors with PSB-0739. Overnight incubation of P4 retinal wholemounts with probenecid or PSB-0739 had dramatic effects on Hmox1 microglial activation across all morphometric parameters. Blocking either purinergic release or sensing, shifted Hmox1 microglia into a significantly more ramified or less activated state. This in short significantly decreased circularity, increased perimeter, increased total skeleton length and number of branch points (Figure 5A-D). These changes in morphology are visually apparent in Figure 5J-L.

**Figure 5:**
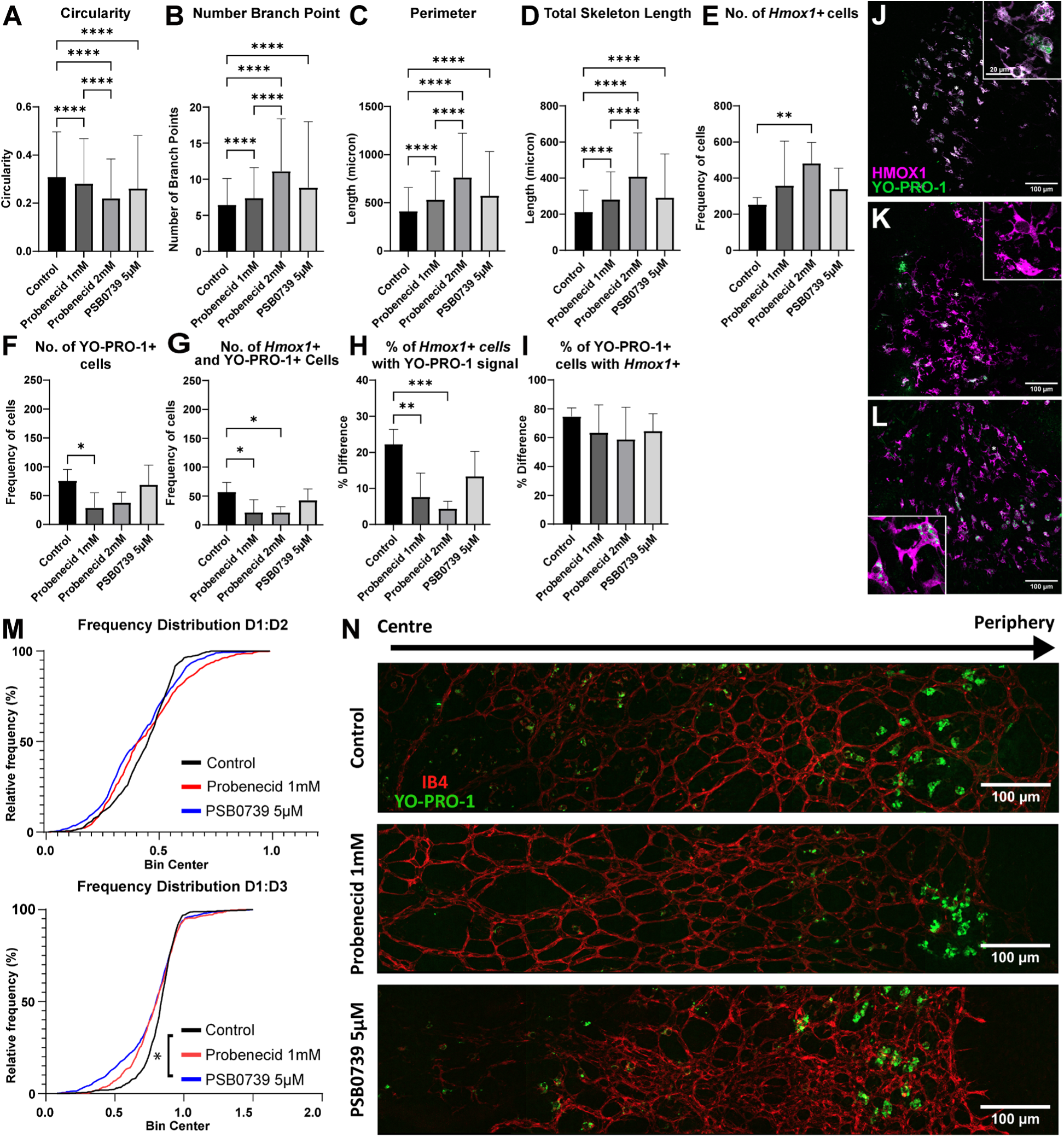
Purinergic signalling through PANX-1 hemichannels and P2Y12 receptors influences Hmox1 microglia activation at P4. Quantification of microglial morphology at P4 under control conditions (black bars) and following pharmacological inhibition using Probenecid 1 mM (darkest grey), Probenecid 2mM (medium grey), and PSB0739 5μM (light grey) for 24 hours. Statistical analysis was conducted using one-way ANOVA followed by Tukey’s multiple comparisons test. Data are presented as mean ± SEM; *p < 0.05, **p < 0.01, ***p < 0.001, ****p < 0.0001. n= 8-10 retinas per group. Cells per group: control= 6829, Probenecid 1mM= 5401, Probenecid 2mM= 4266, and PSB0739 5μM= 3099. **A.** Circularity was significantly reduced in Probenecid 2 mM, and PSB0739 5μM compared to control. **B.** Probenecid and PSB0739 in general significantly increased the number of branch points compared to control. **C.** Cell perimeter was significantly increased by Probenecid 2mM and PSB0739 5μM relative to Control. **D.** Total skeleton length was significantly higher in Probenecid and PSB0739 5μM compared to Control. **E.** Total number of Hmox1 positive cells observed after incubation of probenecid (1mM or 2mM), or PSB0739 5μM for 24 hours prior to fixation and immunolabelling. **F.** Total number of YO-PRO-1 positive cells observed after incubation of probenecid (1mM or 2mM), or PSB0739 5μM for 24 hours prior to fixation and immunolabelling. **G.** Total number of double positive YO-PRO-1 and Hmox1 cells observed after incubation of probenecid (1mM or 2mM), or PSB0739 5μM for 24 hours prior to fixation and immunolabelling. **H.** Percentage of Hmox1+ cells that co-localise with YO-PRO-1 signal, after incubation of probenecid (1mM or 2mM), or PSB0739 5μM for 24 hours prior to fixation and immunolabelling. **I.** Percentage of YO-PRO-1 positive cells that express Hmox1, representing the proportion of apoptotic cells associated with Hmox1 positive microglia after incubation of probenecid (1mM or 2mM), or PSB0739 5μM for 24 hours prior to fixation and immunolabelling **J.** Micrograph displaying overlap of apoptotic cells labelled with YO-PRO-1 and HMOX-1 positive microglia with higher magnification inset indicated by asterisk. **K.** Micrograph displaying reduced overlap of apoptotic cells labelled with YO-PRO-1 and HMOX-1 positive microglia after 24hr incubation with Probenecid 2mM with higher magnification inset indicated by asterisk. **L.** Micrograph displaying reduced overlap of apoptotic cells labelled with YO-PRO-1 and HMOX-1 positive microglia after 24hr incubation with PSB0739 5μM with higher magnification inset indicated by asterisk. Scale bar = 100µm. Inset scale bar = 20µm **M.** Top-Cumulative frequency distributions of apoptotic cells D1/D2 ratio. N=3344. Bottom- Cumulative frequency distributions of apoptotic cells: D1/D3 ratio Key: Control (black), Probenecid-treated (red), PSB0739-treated (blue). N=3344. **N.** Top-Representative image of P4 mouse retinal whole-mounts showing the SVP (IB4, red) and apoptotic cells (YO-PRO-1, green) across the centre-peripheral axis gradient from left (ONH region) to right (vascular leading edge). Middle-Retina incubated in probenecid for 24hrs (1mM). Bottom-Retina incubated in PSB0739 (5μM) for 24hrs. Scale bars = 100μm.

Blockade of purinergic signalling also had a noticeable effect on the numbers of Hmox1 microglia and YO-PRO-1 positive cells. The number of Hmox1 microglia found in the retina increased across the board, although only probenecid 2mM displayed a significant increase (Figure 5E). Unsurprisingly, blocking release or uptake of ATP drastically reduced the number of YO-PRO-1 positive cells, but with probenecid treatment, this may be a mechanistic effect of blocking the PANX-1 channels (Figure 5F). Similarly, the number and percentage of Hmox1 microglia containing apoptotic cell fragments in their vesicles also decreased after blockade. However, this only reached significance with probenecid treatment (Figure 5G, H). Blocking purinergic signalling did not seem to have any effect on the percentage of YO-PRO-1 positive cells, which were colocalised with Hmox1 microglia (Figure 5I).

In addition to morphometrics and cell frequency changes, we also examined the peripherality of YO-PRO-1 positive cells after drug application. To normalise the apoptotic cell locations to account for the changing retina size we converted the origin locations into the D1/2 metrics. These metrics showed no significant changes but there was a trend for probenecid-treated retinas to display a broader apoptotic distribution, with a shift toward the retinal periphery. On the other hand, D1/3 metrics displayed a significant shift of the YO-PRO-1 positive cells treated with PSB-0739 away from the SVP edge and the peripheral non-vascularised areas towards the central retina (Figure 5M, N).

### Stage II Retinal Waves Display Centrifugal Expansion and Modulation by Purinergic Signalling

Because ACCs are present exclusively during the period of Stage II retinal waves, we hypothesised a link between these clusters and wave generation or propagation. To test this, we performed whole-mount calcium imaging using the bath loaded cell permeable dye Calbryte 520AM on IB4-labelled retinas with application of the PANX-1 blocker, probenecid to map wave origination points and quantify wave metrics relative to the vasculature. Representative calcium imaging and wave activity plots are shown in Figure 6A, B. Given the tight anatomical association between ACCs and apoptotic cells expressing PANX-1 hemichannels, we next investigated whether blocking purinergic release altered wave dynamics.

**Figure 6:**
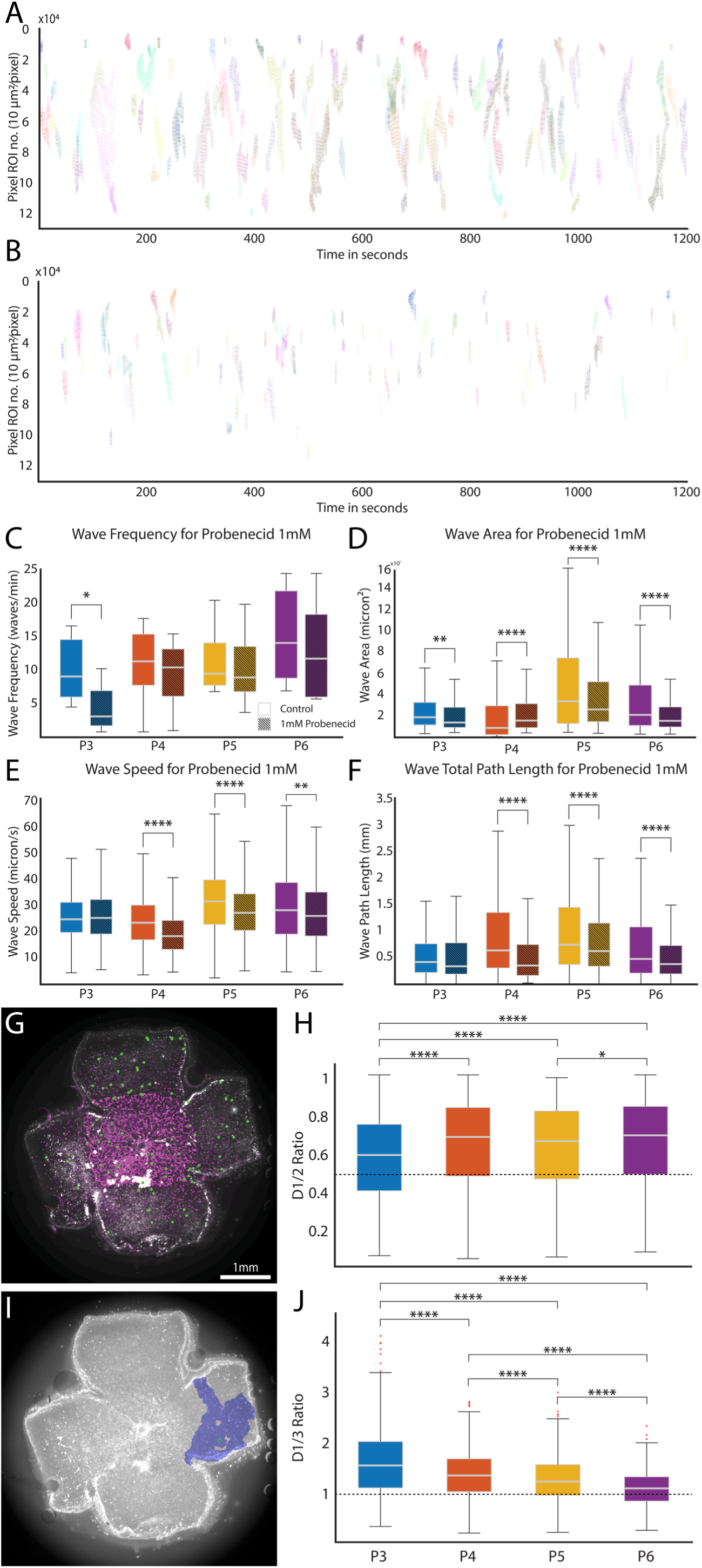
Centrifugal Progression of Retinal Wave Origination is Driven by PANX1-Mediated Purinergic Signalling. **A.** Calcium retinal waves plots of waves in control conditions. Each row represents a pixel ROI (10µm x 10µm) within the calcium imaging recording. Waves are indicated by synchronised activity across channels. The activity is shown against time (in seconds). **B.** Probenecid 1mM profoundly reduces retinal excitability, with significant decrease in wave frequency and number of channels recruited within waves. **C.** Retinal wave frequency increases from P3-P6 while ATP release blockade with probenecid reduces frequency across the board. **D.** Wave retinal area coverage is reduced significantly by probenecid treatment. **E.** Retinal wave propagation speed is reduced by probenecid treatment in the later stage of first postnatal week. **F.** Retina wave path length increases from P3-5 and is significantly reduced by probenecid treatment. Statistical analysis was conducted using Man-Whitney rank sum test. *p < 0.05, **p < 0.01, ***p < 0.001, ****p < 0.0001. **G.** Mouse retinal wholemount (P4) displaying superior vascular plexus (SVP) (labelled with isolectin B4 (IB4)) and the origin points of spontaneous retinal waves recorded with calcium imaging (green spots). **H.** Retinal wave origination becomes increasingly more peripheral (D1/2 > 0.5) over the P3-P6 period. **I.** Example propagation extent of a retinal wave. **J.** Retinal waves originate in the non-vascularised more peripheral regions (D1/3 > 1) of the retina regardless of age.

Treatment with 1 mM probenecid profoundly attenuated nearly all measured wave metrics. Specifically, wave frequency (Figure 6C), speed (Figure 6E), and total path length (Figure 6F) were significantly reduced upon drug application. While probenecid treatment led to a significant decrease in wave area at P3, P5, and P6, P4 retinas uniquely exhibited a significant increase in this metric (Figure 6D).

By aligning functional activity with anatomical landmarks, we mapped all wave origination points relative to the SVP in the P4 retina (Figure 6G) and visualised the spatial coverage of individual waves (Figure 6I). Across all postnatal ages, the majority of retinal waves originated in the peripheral regions. These wave initiation points exhibited a centrifugal expansion pattern mirroring the progression of the ACCs, vasculature, and microglia. To best represent this, we calculated the distance from the optic nerve head to the object (D1) and through to the retinal periphery (D2). The D1/2 ratio metric approaches 1 when the relative distance between the wave origin point is close to the edge retina, regardless of physical size. We also created a version of the ratio which is relative to the SVP border position. Accordingly, a ratio of D1/3 <= 1 defined an object within the vascularised area, while D1/3> 1 was in the non-vascularised area indicated a position in the non-vascularised region (see Figure S2 for details).

Between P3 and P6, wave origins demonstrated a significant increase in relative peripherality (D1/D2; Figure 6H). This peripheral bias was even more pronounced when origins were normalised to the vascular margin; wave initiation was significantly more likely to occur within non-vascularized regions across all stages (Figure 6J). Indeed, the median D1/D3 value, representing the position relative to the vascular front; only approached 1 at P6, when the retina is nearly fully vascularised.

### Synchronised Centrifugal Expansion Pattern Across Multiple Developmental Processes

To examine the overall developmental expansion pattern, we extended the use of the D1/2 metrics across retinal wave generation, vascular growth, apoptotic activity, and microglial positioning. Visualising the relative positions of these processes highlights a strongly conserved organisation (Figure 7). Spontaneous wave initiation points, recorded via MEA (orange) (see Figure S3 for details) or calcium imaging (cyan), predominantly emerge within the non-vascularized periphery and shift further toward the retinal margin between P3 and P6. This centrifugal progression converges at P6 as the SVP achieves near-total coverage. Similarly, the density of apoptotic cells (YO-PRO-1; blue) shifts peripherally over this period. Notably, the expansion of the SVP (magenta) consistently lags behind the wave initiation front, while the Hmox1*+* microglial annulus (red) straddles both the vascular border and the ACC positions. Throughout development, ACCs maintain the most central position relative to these expanding fronts until P6, when they eventually reach the retinal periphery (brown).

**Figure 7.**
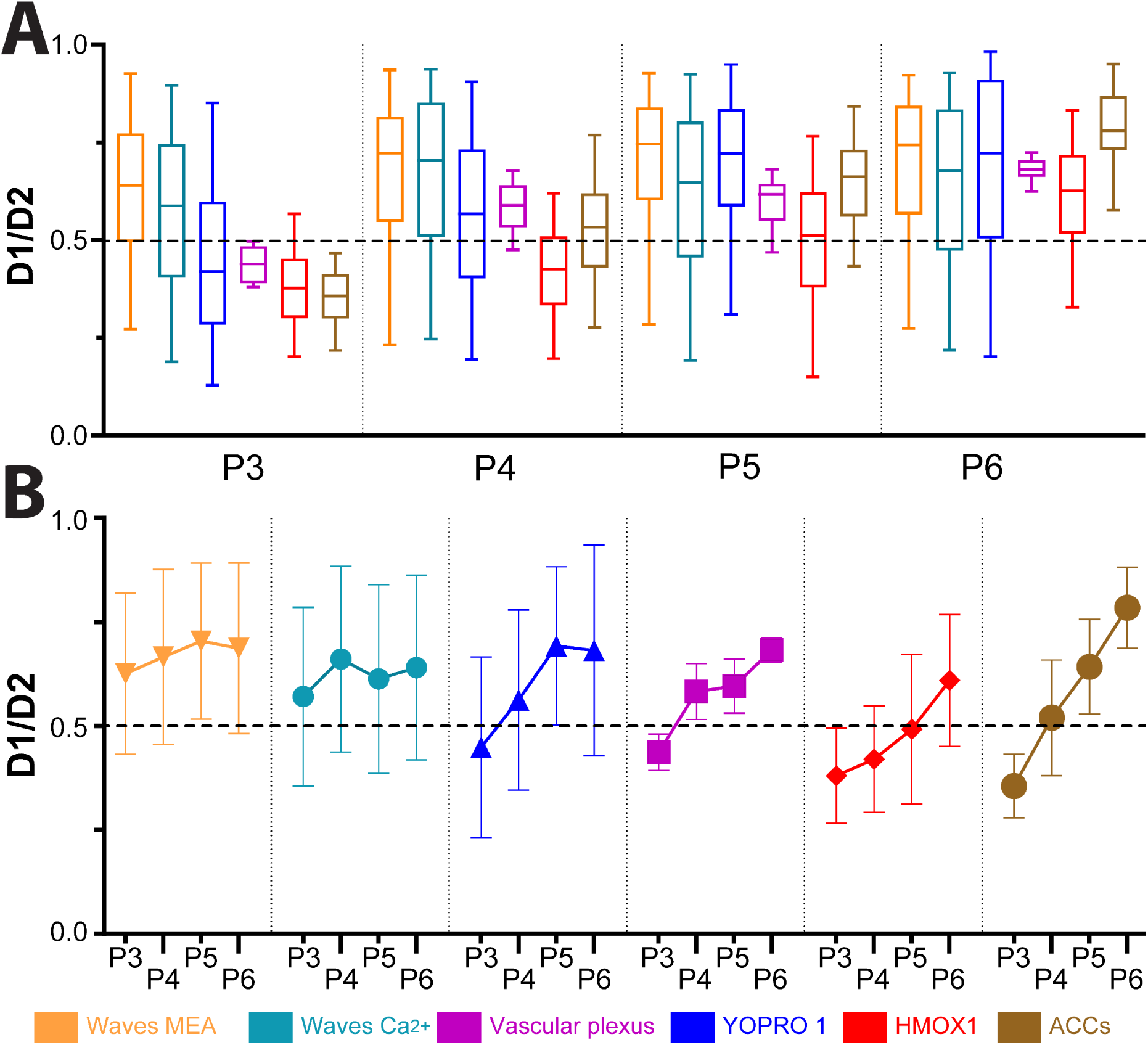
Relative Peripherality of Multiple Developmental Processes Is Age Invariant. **A.** Box plot showing developmental changes in D1/D2 peripherality ratios for stage II retinal waves recorded by MEA, and calcium imaging, SVP extent, YO-PRO-1 labelling, HMOX-1 microglia locations, and ACCs. Each box illustrates the median (horizontal line) and interquartile range, with minimum and maximum values (whiskers). The distribution of each marker of interest increases in peripherality (D1/2 > 0.5) over the P3-P6 period, see Figure S2. **B.** Mean and standard deviation plots for each functional or anatomical measures expressed in terms of D1/2 peripherality metrics across the P3-6 time period

In summary, these data reveal a precisely orchestrated spatiotemporal sequence in the developing retina, where a peripheral wave of neuronal activity and RGC apoptosis consistently precedes vascular expansion. The stable spatial relationship between Hmox1-positive microglia, ACCs, and the trailing vascular front suggests that the outward migration of the SVP is not a stochastic process, but rather one guided by a pre-patterned landscape of purinergic signalling and phagocytic remodelling. This ’wave-front’ architecture ensures that metabolic demand and neuronal maturation are tightly coupled with the emerging blood supply during the first postnatal week.

## Discussion

Herein, we demonstrate for the first time clusters of auto-fluorescent cell complexes which are found in mouse retina over the first postnatal week. Investigating their nature and composition has revealed a tightly regulated synchrony of multiple developmental processes and illuminated several new insights in retinal development and angiogenesis, Figure 8A-C. Dying RGCs in the periphery release purinergic molecules, which help initiate retinal waves and act as a proangiogenic marker, which in turn helps guide the outward expansion of the vascular plexus. This vascular wavefront is coordinated by a distinct microglial annulus that actively monitors the retinal environment Figure 8B. Upon contact, these microglia initiate the phagocytic clearance of apoptotic RGCs, resulting in the formation of ACCs, a process that effectively pre-patterns the retinal landscape for subsequent vascular expansion Figure 8C.

**Figure 8:**
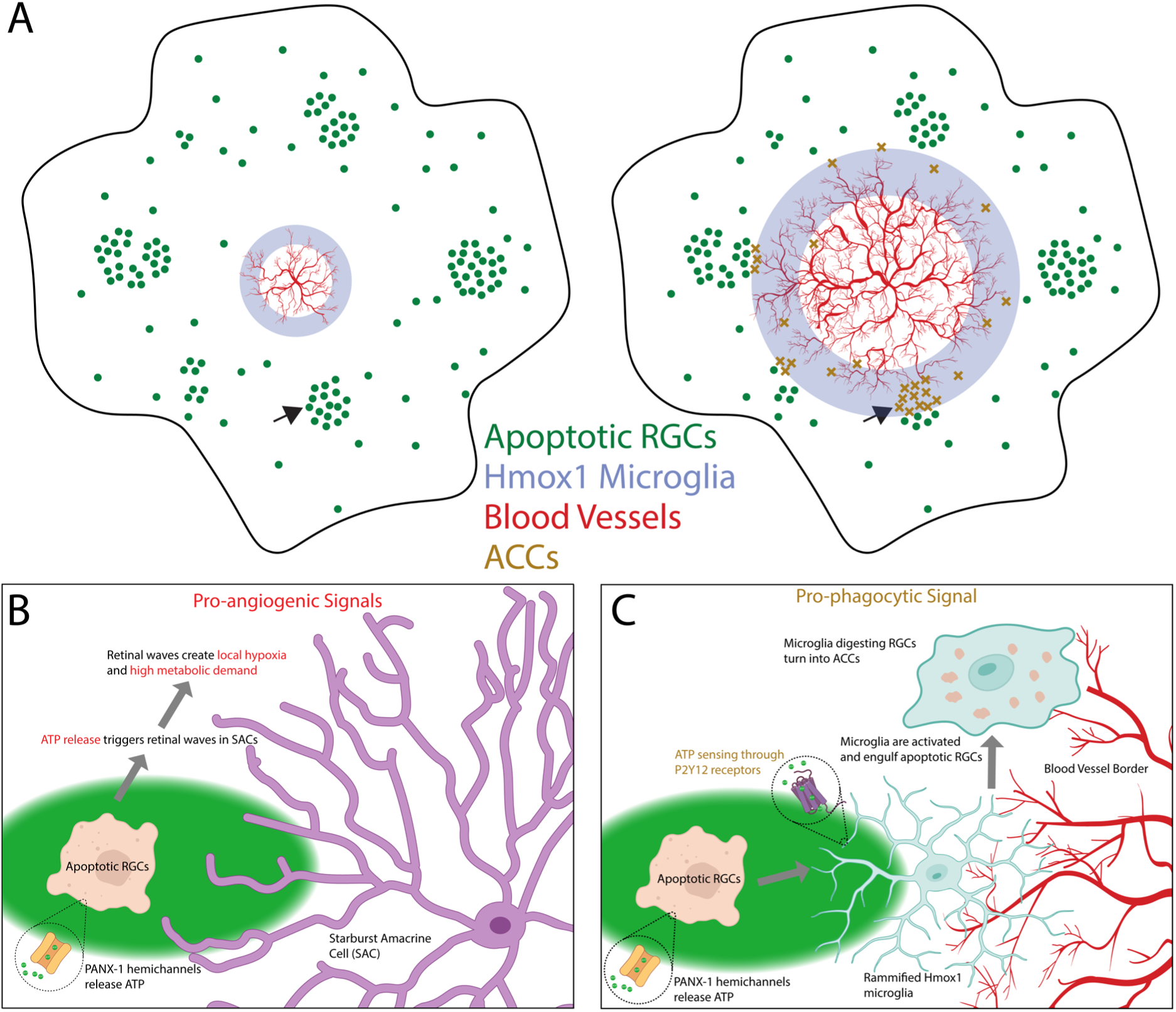
Overview of Synchronised Development in Retina. **A.** Diagram of a retina at two different timepoints displaying the organisation of apoptotic RGCs, the SVP, the annulus of Hmox1 microglia, and ACCs. Arrows indicate areas explored in higher detail in B,C. **B.** Pro-angiogenic and spontaneous activity phase of the apoptotic RGCs releasing purinergic markers through PANX-1 hemichannels. **C.** Phagocytic phase where Hmox1 microglia sense apoptotic RGCs and engulf them for destruction, creating ACCs.

More specifically, we have uncovered a developmental gradient that expands in a centrifugal manner from the ONH. We discovered that apoptosis of developing RGCs promotes the functional upregulation of PANX-1 hemichannels. These dying cells release a host of purinergic molecules such as ATP through the PANX-1 hemichannels. ATP acts to promote the initiation of retinal waves; blocking these PANX-1 channels greatly reduces the frequency and size of the waves. The apoptotic cells and wave initiations are more prevalent in the peripheral non-vascularised zones of the retina. The purinergic release and energy demand of wave initiations, in turn, act as a proangiogenic signal which drives the expansion of the SVP. A specialised subset of microglia expressing *Hmox-1* site astride the SVP edge and help guide the growth of endothelial tip cells. They also have a strong phagocytotic role, where they engulf apoptotic RGCs under the advancing vascularised area. Our findings indicate that microglia engulf apoptotic RGCs and sequester them within phagocytic vacuoles for degradation, thereby forming the ACCs that initially prompted this investigation. This model is further supported by our observation that vascular maturation is significantly more advanced in retinal regions where ACCs are present, suggesting these aggregates serve as a focal point for angiogenesis.

It is surprising that ACCs have remained unidentified until now, given their highly stereotyped progression pattern. However, well-documented autofluorescence from cellular debris and molecules such as lipofuscin (Kennedy et al., 1995; Sparrow, 2007) and drusen (Imamura et al., 2006; Spaide & Curcio, 2010) likely confounded previous attempts at identification. Their relative sparsity may have further led to their misclassification as non-specific staining, underscoring the necessity of pan-retinal visualization for reliable detection. Furthermore, their intrinsic autofluorescence complicates traditional immunohistochemical characterization. By leveraging scRNA-seq, we bypassed these optical limitations to reveal the molecular identity of ACCs as multicellular aggregates comprising microglia, apoptotic RGCs, and endothelial components.

The transcriptomic profile of the largest cell cluster (cluster 0) identifies these microglia as central orchestrators of postnatal retinal development. High expression of canonical markers, including *Aif1*, *P2ry12*, *C1qb*, and *Sall1*, reflects a specialised homeostatic state geared toward neurovascular interaction and phagocytic refinement. Specifically, the co-expression of microglial markers with RGC-specific *Rbpms* (Rodriguez et al., 2014) and apoptotic regulators (*Bax*) provides molecular evidence for the active clearance of RGCs (Edlich et al., 2011; Han et al., 2001). This process is likely driven by a *P2ry12*-mediated purinergic response to apoptotic signals and local hypoxia, with *Aif1* (Iba-1) facilitating the necessary phagocytic cup formation.

Beyond debris clearance, the enrichment of *C1qb* and *Sparc* suggests these cells are actively involved in activity-dependent synaptic tagging and migratory modulation, respectively. Meanwhile, the maintenance of *Sall1* expression indicates that while these microglia are highly active in remodelling, they retain a non-reactive, developmental identity. Collectively, these data support a model where Cluster 0 microglia integrate purinergic sensing and complement-mediated refinement to facilitate the precise spatiotemporal maturation of the retinal landscape.

An interesting finding of this research was that the *Rbpms +* cells did not form a separate cluster within the transcriptomics data. This may be due the fact that phagocytic microglia can encapsulate transcripts originating from neurons and other neural cell types, which are detectable by RNA-seq despite not belonging to a common microglial genetic signature. For example, Solga et al. showed that CNS microglia contain neuronal and oligodendrocyte-specific mRNAs that localise within microglia but are not translated. This research group interpreted that this RNA is acquired through phagocytosis or macropinocytosis of surrounding neural cells (Solga et al., 2015). Similarly, in zebrafish, synapse-engulfing microglia identified in situ display neuronal and synaptic gene expression in single-cell RNA-seq profiles, consistent with engulfed neuronal material contributing to the detected transcriptome (Silva et al., 2021). In line with these observations and given our FACS strategy enriching autofluorescent ACCs rather than intact RGCs, we interpret the widespread *Rbpms* expression across ACC-associated clusters as an expected consequence of microglial engulfment of apoptotic RGCs, rather than evidence for a distinct population of viable RGCs that failed to form a separate cluster.

However, it must be noted that this study was unable to directly compare the transcriptomics of ACC+ cells vs ACC-cells. Due to the rarity of the ACC+ population we focused on collected and analysing a large sample of that data. Head-to-head comparison with ACC-data would greatly inform our understanding of the genetic makeup of the different populations.

A significant outcome of this study is the discovery that Stage II retinal wave initiation follows a highly stereotyped, centrifugal expansion pattern. This observation stands in sharp contrast to the prevailing consensus in the field, which has historically characterised wave initiation as a stochastic or random process (Blankenship & Feller, 2010; Firth et al., 2005; Maccione et al., 2014; Stafford et al., 2009; Zheng et al., 2006). Notably, our data indicate that wave origins are consistently biased toward non-vascularised peripheral regions throughout the first postnatal week. While these findings refuted our initial hypothesis that ACCs serve as the primary source of wave initiation, it reveals a previously unrecognised topographical coordination between neuronal activity and retinal maturation. This spatial bias suggests that the "active" zone of the developing retina is defined by the absence of vasculature, potentially linking wave dynamics to local metabolic or purinergic gradients.

Wave-like activity from the retina propagates to the rest of the visual system and are known to help form the proper retinotopy or world view in both V1 and the superior colliculus (SC) (Ackman et al., 2012; Blankenship & Feller, 2010; Firth et al., 2005). Previous *in vitro* retinal MEA recordings has shown that there is a propagation bias for retinal waves to travel along the nasal-temporal axis (Stafford et al., 2009). This direction bias is maintained through the SC and V1 (Ackman et al., 2012). However, none of these studies noted anything that resembled the centrifugal progression pattern described here. It follows that the centrifugal expansion of wave initiations may aid in the progressive development of the entire visual processing circuit throughout the brain. Focusing the retinal activity origins in concentric annuli ensures that the visual system develops from the central to peripheral retina representations. It follows that this pattern of progression may have a large effect of the development of the retinotopic map in the higher areas but that is beyond the scope if this research.

Several methodological factors likely explain why these developmental mechanisms remained previously undetected. First, our use of high-density Microelectrode Arrays (MEAs) provided superior spatial sampling and comprehensive pan-retinal coverage compared to the sparser configurations used in earlier studies. This was complemented by wide-field calcium imaging, which enabled us to localise wave dynamics with micron-scale precision and directly correlate them with the advancing SVP margin. Furthermore, while previous research often pooled data across the first postnatal week, thereby masking developmental shifts, our daily longitudinal analysis preserved the fine-tuned organisation of wave initiation. By avoiding the loss of temporal resolution inherent in data aggregation, we were able to resolve the centrifugal progression that is otherwise collapsed in cross-sectional or pooled datasets.

A defining feature of this research is the remarkable synchrony observed between neuronal activity, programmed cell death, and angiogenesis. By utilising normalised peripherality metrics (D1/D2), we demonstrated that these processes are not merely concurrent but are strictly organised into a mobile "wavefront" that traverses the retina. Throughout the first postnatal week, retinal wave initiation is consistently sequestered within the non-vascularised periphery, shifting further toward the retinal margin in lockstep with the expanding vasculature.

This advancing front is preceded by an annulus of apoptotic RGCs (YO-PRO-1+), the majority of which reside immediately ahead of the SVP. Positioned precisely at this interface, a specialised subset of Hmox1*+* microglia straddles the leading edge of the vascular plexus. These microglia actively engulf the dying RGCs, sequestering them within intracellular vacuoles to form the ACCs. Our genetic and histological evidence confirms that ACCs are the direct product of this phagocytic interaction, located strategically beneath the growing vascular margin. This highly ordered spatial hierarchy persists until P6, at which point the developmental annuli converge at the retinal periphery, signalling the completion of this primary maturation phase.

Our findings highlighted the importance of purinergic signalling through PANX-1 hemichannels in the integrated development of the retina. Modulation of purinergic signalling has been shown to drastically affect retinal wave dynamics, (Stellwagen et al., 1999) but its’ source was not identified. From our work it seems likely that apoptotic RGCs are a great candidate for this modulation. Purinergic molecules are released through PANX-1 hemichannels, which are found in high levels on mature RGCs (Bao et al., 2004; Dvoriantchikova et al., 2006). Sensing extracellular purinergic markers through P2 receptors is the major molecule responsible for transitions from quiescent to activated phagocytotic microglia (Fontainhas et al., 2011; Velasquez & Eugenin, 2014). However, the differential effects of PANX-1 blockade on wave dynamics across the P3-6 period suggest there may be other interactions along the developmental timeline which bias the effects observed. Blockade in younger animals affected wave frequency significantly with no effect on propagation speed. As the retina develops more the blockade of PANX-1 does trend to reduce frequency but significantly reduce other measures such as total path length and wave propagation speed. This could reflect a reduction in overall PANX-1 expression with aging (Ray et al., 2005) or a shift in the role of PANX-1 signalling from wave generation to wave modulation (Sernagor et al., 2003).

In order to fully understand the developmental synchrony uncovered in this work more direct mechanistic investigation is required. Future work in this area should ulitise chronic knockout lines for apoptosis (BAX-KO) or pannexin-1 to characterize the disruption to the normal develop timeline. In addition, a more fine scaled examination of the Hmox1-positive microglia’s role in guidance of the SVP outgrowth and digestion of apoptotic RGCs into ACCs would be beneficial.

However, the discovery of this integrated mechanism in the retina still raises the compelling possibility that a similar "wavefront" governs development across the entire Central Nervous System (CNS). Auto-fluorescent inclusions within microglia, reminiscent of ACCs, have been observed in the ageing brain (Stillman et al., 2023), while purinergic signalling is already known to modulate spontaneous activity in the developing auditory system (Babola et al., 2021). Our findings suggest a universal developmental logic: the programmed over-proliferation of neurons creates a metabolic and purinergic gradient that directs vascular expansion, fine-tuned by specialised microglial activity.

This theory is bolstered by striking temporal correlations elsewhere in the CNS. In the mouse spinal cord, the peak of apoptosis at E12–E13 aligns precisely with the invasion of vessels into deep tissue layers (Nakao et al., 1988; Yamamoto & Henderson, 1999). Similarly, cortical apoptosis at E14 coincides with vascular infiltration from the subventricular plexus (Chen et al., 2017; Gupta et al., 2021; Kuan et al., 2000). We propose that in these regions, Hmox1-positive microglia may perform the same "double duty" observed in the retina: clearing superfluous apoptotic cells while simultaneously harnessing purinergic signals to guide angiogenesis and synchronise neural activity. Ultimately, our work highlights the necessity of a holistic, multi-system approach to resolve how neural activity and vascular architecture are co-constructed during development.

## Methods

### Characterisation of Auto-fluorescent Cluster Complexes

#### Cell Dissociation

Dissected retinas from P5 C57BL/6 mice were kept in oxygenated artificial cerebrospinal fluid aCSF (118mM NaCl, 25mM NaHCO3, 1mM NaH2PO4, 3mM KCl, 1mM MgCl2, 2mM CaCl2, and 10mM glucose) until they were dissociated using the Neurosphere Dissociation Kit (P) (Miltenyi Biotec GmbH, Teterow, Germany (103-095-943)), in accordance with the manufacturer’s instructions.

#### Fluorescence Activated Cell Sorting (FACS)

FACS polypropylene collection tubes were pre-coated with 20% FBS overnight at 4 °C prior to use. Dissociated cells were suspended in 2% foetal bovine serum (FBS) in the pre-coated tubes. 5% FBS collection buffer was added to a coated FACS collection tube just before FACS flow started. The pre-sorted tube was controlled at 4 ֯C, and the collection tube was controlled at 5 ֯C. Fluorescence-Activated BD Aria Fusion with a 130nm nozzle and 10 PSI pressure was used for cell sorting. Auto-fluorescence typically peaks between approximately 380 and 600nm (ultraviolet to red light). Therefore, a combination of five laser fluorescent detectors was utilised to maximise yield. The auto-fluorescent cell population was sorted first by cells that were auto-fluorescent, then by area, and lastly by width.

#### scRNA-Seq Sequencing

Following FACS, the final cell suspension was centrifuged at 500 x g for 10 mins. All liquid was discarded from the top, leaving 30µl (max volume able to be inserted into the 10x Genomics microfluidic chip). The remaining sample was loaded on the 10x Chromium controller for scRNA-Seq conducted in accordance with the protocols established by 10x Genomics - Universal 3’ Gene Expression (Chromium Next GEM Single Cell 3ʹ Kit v3.1, 4 rxns PN-1000269).

scRNA-Seq was conducted by the Genomic Core Facility and the Bioinformatics Support Unit at Newcastle University, achieving a sequencing depth of 50,000 reads per cell through Illumina NovaSeq 6000. The binary base call (BCL) files underwent de-multiplexing through CellRanger mkfastq version 3.01 to create FASTQ files, followed by alignment and quantification against the human reference genome GRCh38 utilising CellRanger count. Quality control assessments were conducted for each sample in R, eliminating any cells with fewer than 1000 reads, fewer than 500 genes, or more than 10% mitochondrial reads. Additionally, cells expressing haemoglobin genes were excluded from the analysis. The identification of doublets was performed using DoubletFinder, and these were subsequently filtered out from the dataset. Seurat (version 4.3.0) was then used for downstream analysis. The data was normalised and scaled to reduce intercellular technical variation and account for differences in gene expression levels, with nFeature_RNA, percent.mt, and nCount_RNA regressed out during scaling. Principle component analysis was then performed using the 2000 most highly variable genes, followed by graph-based Louvain clustering. Retinal cells were subsequently extracted from the individual samples, and batch effects were mitigated using Harmony (version 0.1.1) to generate a unified dataset. The data visualisation was performed through Uniform Manifold Approximation and Projection (UMAP).

#### Cluster Analysis

Qiagen Ingenuity Pathway Analysis (IPA) software was used for in-depth examination of the differentially expressed gene lists derived from RNA sequencing via IPA core analysis. IPA synthesises meticulously curated information from a variety of biological data sources, including gene expression, molecular interactions, and pathway data from a vast experimental database. This synthesis facilitates the identification of crucial biological signals. The outcomes of the differential expression analysis were refined to retain only those with a p_val_adjust of 0.05 or lower for each cell type, subsequently uploaded to IPA. The gene symbols were designated as ID with the selection of the ’Gene symbol’ option. The avg_log2FC and p_val_adj were utilised as the ’Observations’, with ’Exprs Fold Change’ and ’Exprs False Discovery Rate’ options applied respectively. Next, the core analysis was conducted by selecting the ’Expression Analysis’ option, with ’Exprs Fold Change’ chosen to compute the z-score. This analysis was performed utilising the ’Ingenuity knowledge base (genes only)’, taking into account both ’Direct and indirect’ relationships, and ’Causal networks’ were selected. Following the completion of the core analysis, the ’Upstream regulators’ option was selected, and the results were organised by z-score to identify the networks exhibiting the highest activation scores.

### Electrophysiology Methods

Retinas were isolated from mouse pups at P2 (N=4 retinas), P3 (N=4), P4 (N=10), P5 (N=14), P6 (N=3), P7 (N=2), P8 (N=2), P9 (N=2), P10 (N=2), P11 (N=2), P12 (N=1), P13 (N=1). The isolated retina was placed, RGC layer facing down, onto the MEA and maintained stable by placing a small piece of polyester membrane filter (Sterlitech, Kent, WA, USA) on the retina followed by a bespoke anchor. The retina was kept in constant darkness at 32°C with an in-line heater (Warner Instruments, Hamden, CT, USA) and continuously perfused using a peristaltic pump (∼1 ml/min) with artificial cerebrospinal fluid equilibrated with 95% O2 and 5% CO2. Retinas rested in-situ for 2 hours before recording, allowing sufficient time for spontaneous activity to reach steady-state levels. Probenecid (4-(dipropylsulfamoyl)benzoic acid) (water soluble form, ThermoFisher Scientific, P36400) was used at 1mM.

High density MEA extracellular recordings of spontaneous waves were performed as described in detail in Maccione et al. (2014), using the BioCam4096 platform with APS MEA chips type HD-MEA Stimulo (3Brain GmbH, Switzerland), providing 4096 square microelectrodes of 21 µm x 21 µm in size on an active area of 5.12 x 5.12 mm^2^, with an electrode pitch of 81 µm. Two P5 and one P4 datasets were acquired with the MEA chip HD-MEA Arena (2.67x2.67mm^2^ active area, electrode pitch 42 µm) for electrical imaging.

Raw signals were visualised and recorded at 7 kHz sampling rate with BrainWaveX (3Brain GmbH, Switzerland). Each dataset consisted of 30 minutes of continuous recording of retinal waves. Following the recording session, retinas were photographed on the MEA to document their precise orientation relative to the electrode array.

### Calcium Imaging Methods

#### Tissue Preparation

C57BL/6 pups from P3 (N=8), P4 (N=15), P5 (N=11), and P6 (N=8) were enucleated, and retinas were dissected at room temperature in oxygenated ACSF under a dissecting microscope. Isolated retinas were mounted whole onto a hydrophilic PTFE membrane (H100A047A Advantec, Japan) with the retinal ganglion cell layer facing upwards. All further incubations were conducted at 35C° in a hyperoxygenated darkened chamber. P4 and older retinas underwent incubation in 0.0001% Pronase E (P5147 Sigma-Aldrich) for 30 minutes to permeabilise the inner limiting membrane (ILM) (Dalkara et al., 2009). To label the extent of vascularisation, the retinas were incubated in isolectin B4 594 (DL-1208, Vector Labs) for 30 minutes at 10 µg/ml (Stucky & Lewin, 1999). Retinas were bath loaded with the calcium indicator Cal 520 AM (AAT Bioquest) (10µM) for 2 hours and were then flattened further with a second piece of PTFE membrane placed on top and before transferring to the imaging microscope. The retinas were perfused with fresh oxygenated aCSF with a peristaltic pump at 1.7 ml/min. Tissue was allowed to settle in the perfused heated solution for at least 30 minutes before imaging.

#### Data Acquisition

Imaging was conducted in the Newcastle University Functional and Light-Activatable Multi-dimensional Electrophysiology facility (FLAME). This was conducted on a Nikon FN1 microscope at x4 NA 0.2 (Nikon CFI Plan Apochromat Lambda D 4X) magnification with a x0.63 relay lens resulting in a field of view of 5.3x 5.3 mm and a resolution of 2.6µm per pixel. To enable multiple wavelength imaging the microscope used a quad band Chroma 89402 filter cube. Images were acquired at 470nm and 590nm for the calcium signal and blood vessels respectively. Time series images were acquired at 2Hz for 10 minutes at 2048 x 2048 pixels at 16bit resolution. For drug applications, an appropriate wash in and wash out time was used according to the literature. Probenecid (1mM) was used to block purinergic release from apoptotic cells and examine the effect on retinal wave generation/spread.

#### Data Processing and Analysis

All data processing was conducted in MATLAB. Image time series were motion corrected using 2D cross-correlation approach. To properly visualise the retinal waves the time series were downsampled to 512 x 512 pixels and went through a pixel-wise ΔF/F procedure. The calcium activity baseline was calculated by creating a low-pass percentage filtered trace. Each movie frame (F) was normalized by dividing its difference from the baseline frame (F-F0) by the baseline frame ((F-F0)/F0) to produce a ΔF/F movie.

To track the spatiotemporal properties of the calcium waves the ΔF/F movies were thresholded, and the waves were identified with a semi-automated algorithm to determine their identity. Once this was done, a large variety of metrics were extracted including initiation point, speed, area, frequency, and spatial relation to vascularised area.

All code relating to this project can be found here: https://github.com/GrimmSnark/paperCodeSavage2026

### Immunohistochemistry

#### Immunostaining

Whole-mount retinas were prepared from mouse pups aged P2-P11, flattened on nitrocellulose membrane filters or PTFE membrane and fixed for 45 min in 4% PFA. Some retinas were labelled for apoptotic cells with the selective YO-PRO-1 dye (Y3603, Invitrogen) at 200uM for 1 hour in carbogenated aCSF before fixation. Retinas were incubated with the primary antibodies (RBPMS 1830-RBPMS, rabbit polyclonal, Phosphosolutions (1:500), ChAT AB144P, goat polyclonal, Merck Millipore (1:500),VChAT PA5-77386, rabbit polyclonal, ThermoFisher Scientific) (1:500), Iba-1 019-19741, rabbit polyclonal, Alpha Labs (1:1000), HMOX-1 AB_2118685, rabbit polyclonal, Proteintech (1:1000), PANX-1 488100, rabbit polyclonal, Invitrogen (1:500)) with 0.5% Triton X-100 in PBS for 3 days at 4°C, then washed with PBS and incubated with the secondary antibody solution for 1 day at 4°C (with or without IB4). Secondary antibodies included 0.5% Triton X-100 with donkey anti rabbit Alexa 568 (1:500), donkey anti-goat Dylight 488 (1:500), goat anti-rabbit A546 (1:500). Isolectin B4 (IB4) staining of blood vessels (1:250) was performed together with the secondary antibody staining step. Finally, retinas were washed with PBS and embedded with Vectashield (H-1900, Vector Laboratories).

For a specific antibody lot of PANX-1, the tissue underwent an additional antigen retrieval step in a water bath at 80°C for 30 minutes using sodium citrate. Retinas were then incubated in blocking solution (5% secondary antibody host species serum with 0.5% Triton X-100 in PBS) for 1 hour.

#### Imaging

Zeiss AxioImager with Apotome and a Zeiss LSM 800 confocal microscope were used to image the retinas. High-resolution images of the RGC layer down to the inner nuclear layer (INL) were obtained by subdividing retinal wholemounts into adjacent smaller images that were subsequently stitched back together to view the entire retinal surface. Regions of interest were selected around the clusters.

To compensate for variability in retinal thickness, several focus points were set across the retinal surface to maintain sharp focus on the desired cell layer. Each picture was then acquired in all colour channels at 20x magnification, and with 10% overlap between neighbouring areas. This overlap is used to correctly align and stitch together all pictures using the Zen Pro software (Zeiss). Some wide-field fluorescent images were acquired with the Nikon Ni-E microscope. Grids of 5x5-8x8 images were taken at 10x or 20x, and then combined to obtain a high-resolution image of the whole retina

#### Peripherality and Vascular Relation Metrics (D1/2, and D1/3)

To determine the relative position of objects—including ACCs, labelled cells, and wave origins, we calculated the distance from the optic nerve head to the object (D1) and through to the retinal periphery (D2). The ratio (D1/D2) provides a measure of an object’s relative peripherality, independent of individual retinal size. Additionally, for a subset of samples, we measured the distance from the optic nerve head through the object to the margin of the superficial vascular plexus (D3). Accordingly, a ratio of D1/3 <= 1 defined an object within the vascularised area, while D1/3> 1 was in the non-vascularised area indicated a position in the non-vascularised region (see **Figure S2** for details).

For the ACC D1/2 values (Figure 1F) one-way ANOVA was used on all 233 ratio values for all eight groups. Tukey post-hoc test was used to identify significant changes in cluster positions between consecutive developmental days.

#### Vascular Plexus Analysis

Sholl analysis was performed using FIJI to quantify blood vessel branches. IB4 stained vasculature images from retinal wholemounts were rendered binary using the ‘Threshold’ function. To remove signals inherent to auto-fluorescence of the ACCs, a separate colour channel image showing only cluster cells was rendered binary and subtracted from the vasculature image. Subtracted images were then adjusted using the ‘Brightness/Contrast’ function to obtain solid binary images.

To quantify blood vessel density in the vicinity of ACCs, segmental sections of the binary, cluster-subtracted whole retina vasculature images were outlined using the ‘Angle’ tool (allowing measurement of the angle of the segment and subsequent calculation of the surface area of the region of interest (ROI)) either with or without a cluster at the edge of the vascular plexus. Sholl rings were set from the centre of the ONH with a step size of 30 µm. Measurements were not taken from the inner 50 % of the segment (based on the radius of the vascular plexus at its largest point). Intersections were counted manually.

To quantify blood vessel branch density in areas with ACCs versus areas without at matching eccentricities, ROIs were outlined using the ‘Polygons’ function either around clusters (ACC+ ROI), or in adjacent ACC negative regions at similar eccentricities (ACC-ROI). This was done in images from a colour channel in which the blood vessels were not visible. Once all ROIs were outlined for one retina, they were overlaid on the binary, cluster subtracted vasculature image for that same retina. Any ROIs noticeably outside of the vascular plexus were discounted. Sholl rings were set from the centre of the ONH with a step size of 30 µm. Intersections were counted manually.

All cell counts were performed using FIJI. Cluster cells were counted manually in retinal wholemount images using the ‘Multi-point’ function. Cells were identified by auto-fluorescence using an image captured on a channel without any fluorescing immunostaining. The vascular plexus was outlined using the ‘Freehand’ tool and ‘Measure’ function was used to determine surface area. Clusters were delimited and cells counted manually, identified by auto-fluorescence. A ‘cluster’ was defined as an uninterrupted group of auto-fluorescent cells spatially separated from any other group, localised near the outer edge of the superficial vascular plexus. No lower or upper size limit was assigned to clusters.

#### ACCs and Microglial Densities

To quantify microglia and ACC densities, radial straight lines were drawn from the centre of the ONH to the periphery of the retina, either through the centre of an ACC group or through an adjacent ACC negative region. This was only performed where radial lines reached the periphery of the retina, uninterrupted by cuts. ROIs were outlined using the ‘Oval’ selection tool at four points along each radial line to comprise an ROI set. The four ROI types in each set were ahead of the ACC positive/negative region, at the ACC positive/negative region, behind the ACC positive/negative region, and close to the ONH. ROIs were outlined at comparable eccentricities for each set. Iba-1 stained microglia were counted manually. All Iba-1 stained whole retinas were from P5 or P6 animals. ACCs were identified by auto-fluorescence and counted manually.

#### Quantification of microglia, apoptotic cells and co-localisation

For retinal wholemounts stained for HMOX-1 and YO-PRO-1 binary masks were created with FIJI/ImageJ, Labkit (Arzt, M. et al 2022), and custom MATLAB code. Labelled cells were counted and co-localisation percentages were also calculated. This was based on the HMOX-1 microglia mask containing any positively labelled pixels from the YO-PRO-1 channel. Statistical testing between different drug conditions was conducted with a Kruskal-Wallis ANOVA test with multiple comparisons, threshold at p= 0.05.

### Microglia Morphometry

#### Computational Microglia Labelling

Images were processed and filtered using FIJI/ImageJ (Schindelin et al., 2012). Images were cleaned for background noise using Gaussian convoluted background subtraction (σ = 50 pixels) and median filtering (3 x 3 pixels). Single colour image channels depicting stained cells of interest (YO-PRO-1, Hmox1 etc) were segmented with Labkit (Arzt, M. et al 2022) and custom code (https://github.com/GrimmSnark/paperCodeSavage2026). Quantitative measurements of the labelled microglia were conducted utilising the FIJI plugin “Measure Microglia Morphometry” (Martinez et al. 2023). Briefly, this plugin quantifies a plethora of morphological metrics including area, perimeter, convex area, average branch length, and branch point number (Figure S1.)

#### Microglial Behavioural Parameters Statistical Analysis

Statistical analysis and data visualisation were performed using GraphPad Prism 10. Two-tailed Mann-Whitney tests and non-parametric ANOVA were applied, with a significance threshold set at P ≤ 0.05. Given the non-parametric nature of the data distribution, results were presented as the median accompanied by a 95% Confidence Interval (CI). The significance levels are indicated as follows: not significant (ns) for P > 0.05, * for P ≤ 0.05, ** for P ≤ 0.01, *** for P ≤ 0.001, and **** for P ≤ 0.0001.

## Supporting information

Supplementary Table 1 References

## Acknowledgements

We are grateful to Leverhulme Trust (RPG-2022-061), BBSRC (BB/T017627/1) and EPSRC/ERC (EP/Y031016/1) for funding this work. We would also like to thank the Newcastle University FMS Bioimaging Unit for their continual support during this project. For the purpose of open access, the author has applied a Creative Commons Attribution (CC BY) licence to any Author Accepted Manuscript version arising from this submission.

This work is dedicated to the memory of Evelyne Sernagor, who conceptualised and led this study until her passing in March 2025. Her brilliance and mentorship remain the foundation of this research.

Authour contributions (CRediT):

**Michael A. Savage:** Conceptualization, Methodology, Software, Validation, Formal analysis, Investigation, Data Curation, Writing - Original Draft, Visualization, Project administration

**Cori Bertram:** Methodology, Validation, Formal analysis, Investigation, Data Curation, Visualization

**Jean de Montigny:** Conceptualization, Software, Formal analysis, Investigation, Data Curation, Visualization

**Courtney A. Thorne:** Investigation, Formal analysis, Visualization

**Rachel Queen:** Software, Formal analysis, Visualization

**Majlinda Lako:** Conceptualization, Methodology, Resources, Supervision, Writing - Review & Editing, Project administration, Funding acquisition

**Gerrit Hilgen:** Writing - Review & Editing, Supervision

**Evelyne Sernagor:** Conceptualization, Methodology, Formal analysis, Investigation, Resources, Writing - Original Draft, Supervision, Project administration, Funding acquisition

## Supplementary Info

**Figure S1:**
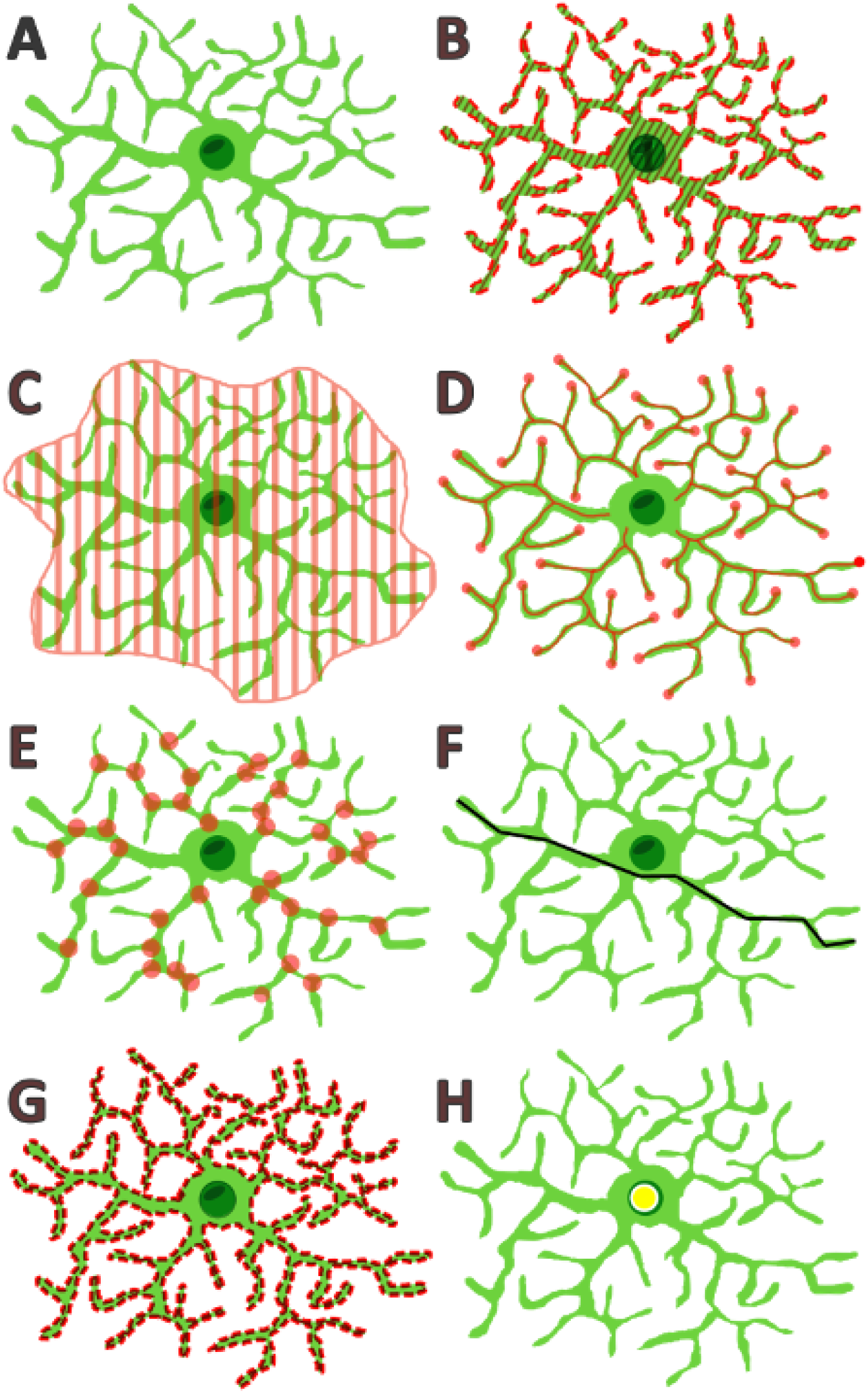
Microglial Morphometric Parameter Measurements. **A.** Microglial cell shape example, **B.** Area of the cell is illustrated by red stripes covering the cell. **C.** Convex Area illustrated by area covered in red stripes, **D.** Length of individual branches-shown by a red line and red dot at the end point of each measurement, **E.** Number of branch points shown by a red dot covering every branch intersection, **F.** Geodesic diameter highlighted by a black line along the shortest path through the maximum distance of 2 points of the microglia cell, **G.** Perimeter of the cell show by a dashed red line around the cell outline, **H.** The geometric centre of the cell is represented by a yellow circle.

**Figure S2:**
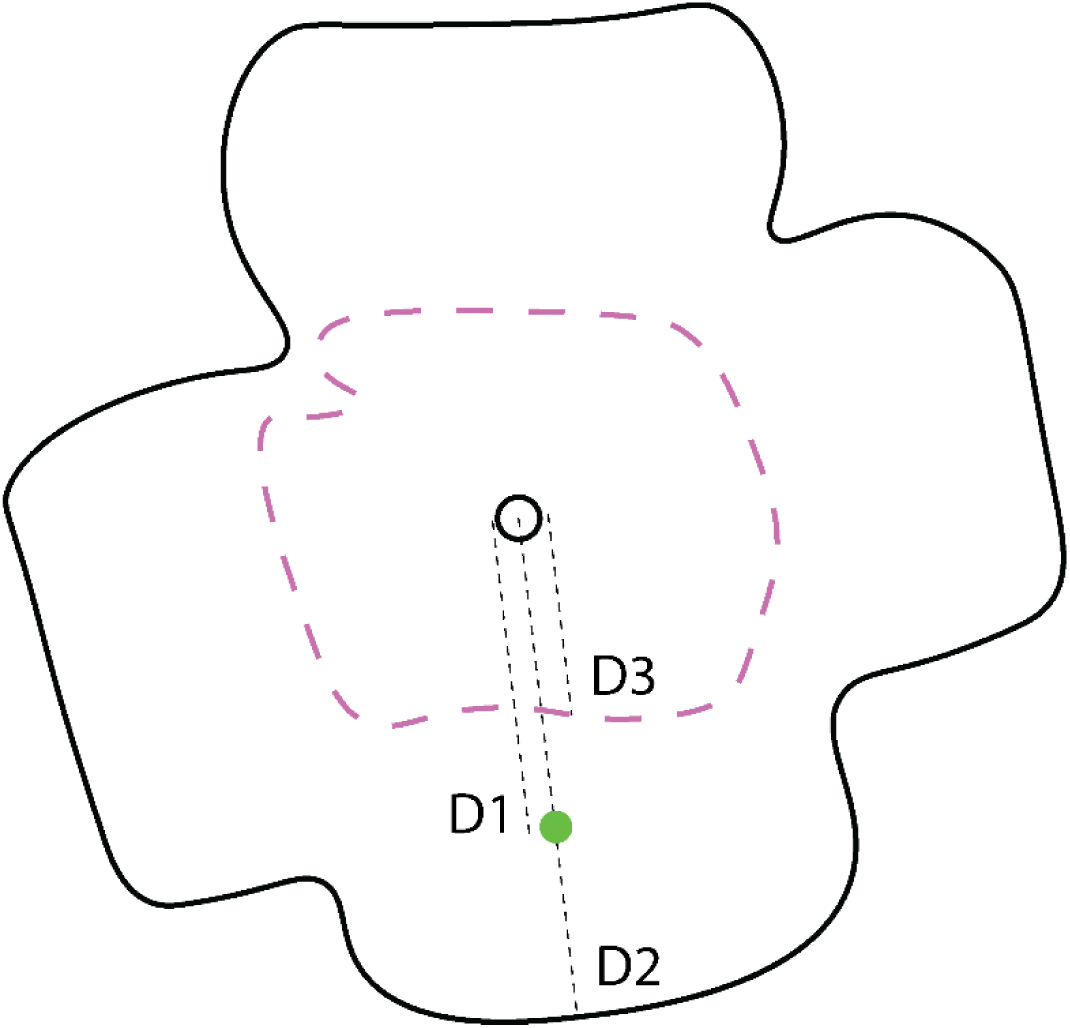
Peripherality and Vascular Relation Metrics. Method for calculating the relative position of anatomical markers between the ONH (small black circle in the middle) the periphery, and the vascular plexus. Anatomical markers such as ACCs are represented by a green dot. D1: distance from centre of ONH to the object of interest. D2: distance between the object of interest to periphery. D3: The distance from the ONH through the line of the object of interest to the vascularised border. D1/2 value greater that 0.5 equates to the periphery of the retina. D1/3 value greater than 1 equates to the non-vascularised area of the retina.

**Figure S3:**
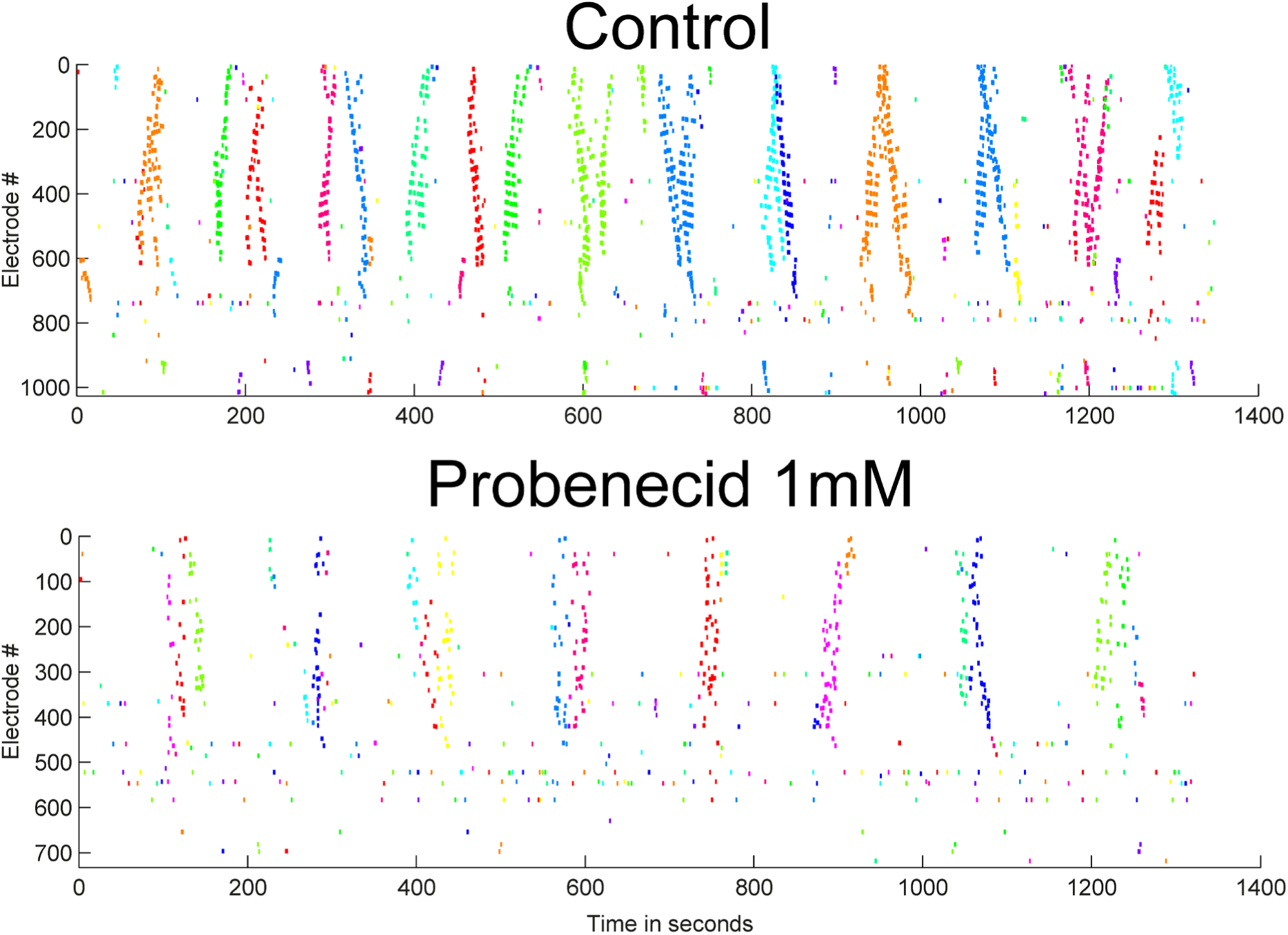
Multi-Electrode Array Recording of Spontaneous Waves Affected by PANX-1 Blockade. Raster plots of waves from a P5 mouse in control conditions, and in the presence of probenecid 1mM (recording started 30 minutes after adding the drug). Each row represents an active electrode on the MEA. Waves are indicated by synchronized activity across channels. The activity is shown against time (in seconds). Waves are detected according to methods described in Maccione et al., 2014. Each detected wave is in a different color. Probenecid profoundly reduces retinal excitability, with significant decrease in wave frequency and number of channels recruited within waves.

## Notes

### Competing Interest Statement

The authors have declared no competing interest.

### Summary of Updates

We have addressed a number of suggestions based on our first submission to eLife.

